# DISTEMA: distance map-based estimation of single protein model accuracy with attentive 2D convolutional neural network

**DOI:** 10.1101/2021.03.29.437573

**Authors:** Xiao Chen, Jianling Cheng

## Abstract

**Background:** Estimation of the accuracy (quality) of protein structural models is important for both prediction and use of protein structural models. Deep learning methods have been used to integrate protein structure features to predict the quality of protein models. Inter-residue distances are key information for predicting protein’s tertiary structures and therefore have good potentials to predict the quality of protein structural models. However, few methods have been developed to fully take advantage of predicted inter-residue distance maps to estimate the accuracy of a single protein structural model.

**Result:** We developed an attentive 2D convolutional neural network (CNN) with channel-wise attention to take only a raw difference map between the inter-residue distance map calculated from a single protein model and the distance map predicted from the protein sequence as input to predict the quality of the model. The network comprises multiple convolutional layers, batch normalization layers, dense layers, and Squeeze-and-Excitation blocks with attention to automatically extract features relevant to protein model quality from the raw input without using any expert-curated features. We evaluated DISTEMA’s capability of selecting the best models for CASP13 targets in terms of ranking loss of GDT-TS score. The ranking loss of DISTEMA is 0.079, lower than several state-of-the-art single-model quality assessment methods. The work demonstrates that using raw inter-residue distance information alone with deep learning can predict the quality of protein structural models reasonably well.

## Introduction

Estimation of protein model accuracy (EMA) or assessment of protein model quality (QA) is an important problem in protein structure prediction. Since the seventh Critical Assessment of Techniques for Protein Structure Prediction (CASP7) [1] EMA (or QA) has been a prediction category in CASP experiments. A lot of methods have been developed to evaluate the quality of protein models [2], [3], [4], [5], [6]. These EMA methods fell into two main categories: multi-model methods and single-model methods. A multi-model method takes a pool of prediction structure models of the same target as input to evaluate their quality based on the similarity between the models and possibly other structural features. A single-model method predicts the quality of a single protein without comparing it to any other structure models. A multi-model method’s performance depends on the proportion of good models in the pool and may perform poorly when there are only a few good models. In contrast, a single-model EMA method [7] can estimate the accuracy of a single protein model without being influenced by the existence of other models. A recent study [8] shows the single-model methods can perform better than multi-model methods in some cases. Moreover, different from multi-model methods that can only predict relative quality of models in a pool, single-model methods can predict the absolute quality of a single model, which is important for users to decide how to use the model. Therefore, single-model quality assessment is receiving more and more attention, even though its average performance was still lower than multi-model methods in the past several CASP experiments.

Numerous machine-learning methods have been developed to combine various protein structural features to assess the quality of protein models. ProQ2 [9] and Model Evaluator [7] applied support vector machines (SVM) with residue contacts, secondary structure information, solvent accessible surface area, and/or sequence features to predict a global quality score – the global similarity between a protein model and its native structure. ProQ3 [9] added the Talaris energy as a new feature on top of the ProQ2. ProQ3D [10] used a multi-layer perceptron with the same features used in ProQ3 for protein model quality prediction. Recently, deep learning-based models have been applied to improve the estimation of model accuracy. DeepQA [3] utilized deep belief networks to predict the global quality score. ProQ4 [11] exploited the transfer learning and 1D convolutional neural network (CNN) to predict the Local Distance Difference Test (LDDT) score [12]. DeepRank [5] applied deep learning to integrate multiple features including residue-residue contact features to predict model quality and performed best in selecting best protein models in the CASP13 experiment. DeepRank2 [6] added a new inter-residue distance feature with a deeper and wider neural network to predict global model quality. Some recent methods leverage more complex deep learning architectures. Treating a protein structural model as a graph, ProteinGCN [13], GraphQA [14] and VoroCNN [15] applied graph convolutional networks (GCN) to estimate the model accuracy. ResNetQA [16] and DeepAccNet [17] used deep residue networks to address the problem.

In addition to the inference technology, the performance of EMA method depends on input features. In CSAP13, DeepRank [5] demonstrated that accurate residue-residue contacts (a simplified representation of distances between residues) predicted by deep learning improved the prediction of the quality of protein structural models, suggesting that more detailed residue-residue distance predictions could further improve EMA. However, only a few methods [6], [16], [17], use residue-residue distances to estimate the accuracy of protein structural models.

Instead of extracting features from the predicted residue-residue distance maps based on human intuition or expertise as most existing methods did, we designed a 2D convolutional neural network (2D-CNN) with the channel-wise attention to directly use the raw difference map between the distance map of a model and the distance map predicted from the protein sequence to estimate the accuracy of a single protein model. On the CASP13 dataset, our method - DISTEMA – achieved the better performance than other state-of-art single-model methods in terms of the ranking loss of selecting the best models for protein targets. The results show that the attentive 2D-CNN methods can automatically extract useful information from raw residue-residue distance maps alone to predict the quality of a single protein model without using other protein structural features.

## Methods and Materials

### Difference Map as Input Feature

We applied a real-value distance predictor DeepDist [18] to predict an inter-residue distance map from the sequence of a protein target as matrix *A* (*L* × *L*), where *L* donates the sequence length and *A* [*i,j*] is the distance between residues *i* and *j*. A was compared with the distance matrix *B* (*L* × *L*) calculated from the coordinates of residues in a protein structure model to generate a difference map *D*. Because *A* can be considered the expected distances between residues and *B* the actual distances between residues in a model, *D* measures how well the model meets the expectation and therefore provides useful information about the quality of the model. Considering that large distances tend to have little impact on the fold of a protein structure, before the generation of the distance map, a distance threshold (i.e., 16 Angstrom) was applied *A* and *B* to filter out the distances that are greater than the threshold. That is, if either *A*[*i*,*j*] or *B*[*i*,*j*] is greater than 16, both *A*[*i*,*j*] or *B*[*i*,*j*] were set to 0, producing two filtered distance matrices *A*\* and *B*\*. The difference map *D* was an element-wise subtraction between *A*\* and *B*\*. Since *A*\* and *B*\* are symmetrical, *D* is also symmetrical. To speed up the training of the deep learning method, the distances in the lower triangle of *D* is set to 0 to produce a matrix *U*. *U* that only contains the values of the upper triangle of *D* is used as input for the deep learning method to predict model quality. For example, **Figure 1** visualizes *A*\*, *B*\*, *D*, and *U* of a model of CASP13 target T0949.

**Figure 1.**
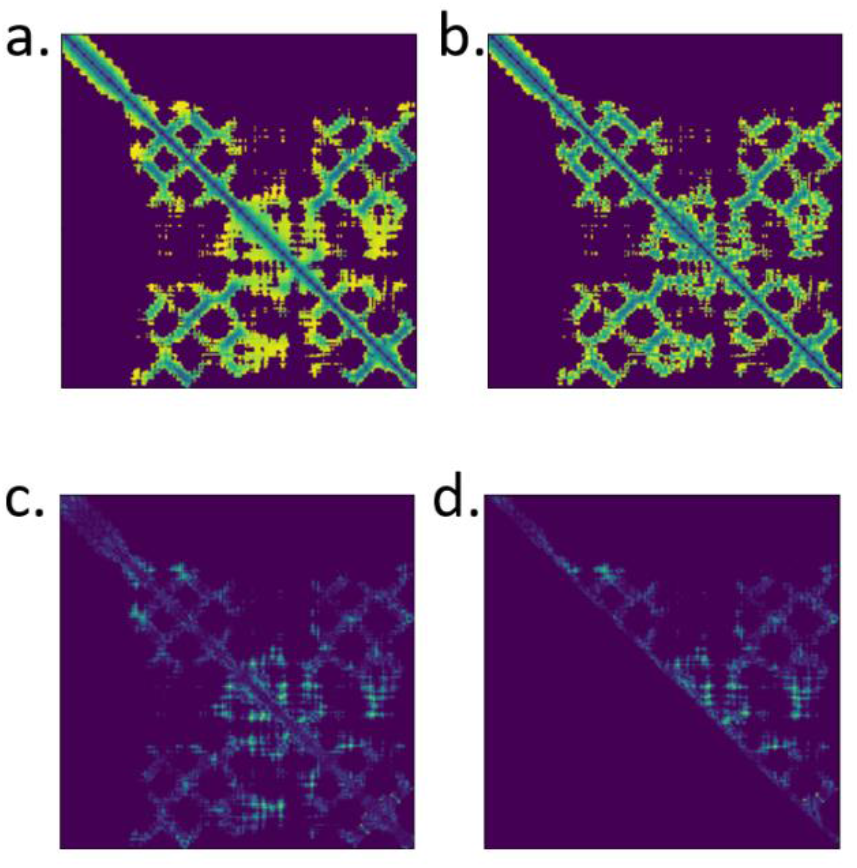
The difference map for CASP13 target T0949. *a* and *b* are filtered matrices from predicted distance map, and model structure distance map. *c* is the element-wise subtraction between *a* and *b*. *d* is the upper triangular part of *c*. *d* is the difference map. For those four maps, the lighter area represents for larger value.

### Deep learning architecture and training

The architecture of the deep learning network of DISTEMA is illustrated in **Figure 2**. The network takes the difference map *U* of a model as input to predict the global distance test total score (GDT-TS) [19] of the model. A GDT-TS score ranging from 0 to 1 measure the quality of a model – the global similarity between the model and is native structure. Higher the score, better is the model quality. A true GDT-TS score of a model can be calculated by comparing a model with its native structure using some tools like TM-score [20] if the latter is known. Otherwise, the GDT-TS score needs to be estimated or predicted from the features of the model.

**Figure 2.**
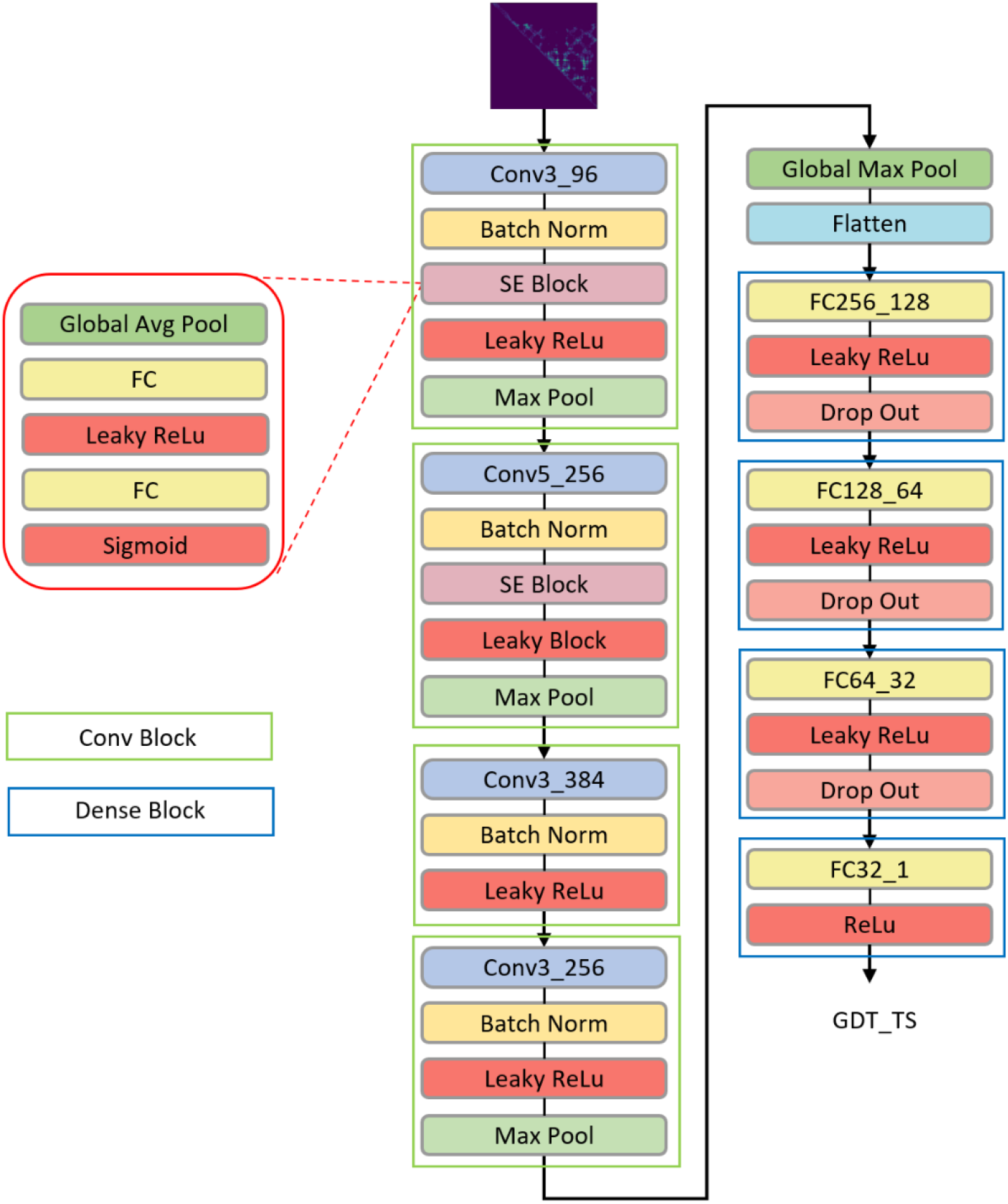
The schematic shows the architecture of DISTEMA. We applied four CONV blocks, two SE blocks, four dense blocks to build this model. The input is difference map and predicted GDT TS score is output.

The protein models of CASP8-12 whose true GDT-TS scores are known were used to train the deep learning method to predict their GDT-TS scores. The input size for the deep learning method is *b* × 1 × *L* × *L*, and the output size is *b* × 1. Here *b* denotes the batch size, 1 × *L* × *L* the size of the difference map, and 1 the number of input channel (i.e., the distance difference value). We let the input in the same batch have the same *L* to speed up training, even though the *Ls* in different batches can be different. The deep network is composed of four convolutional (Conv) blocks, a global pooling layer, a flatten layer, and four dense blocks. The four Conv blocks extract features from the input. Each of the first two Conv blocks contains a squeeze-and-excitation (SE) block [21] with the channel-wise attention mechanism to automatically assign higher weights to more relevant features. Both SE blocks use the same squeeze-and-excitation ratio (i.e., 16). In each SE block, the global average pooling layer extracts single average value from the previous convolutional layer’s channels; two fully connected layers shrank the inner neural size first and then increase the size to the original number; the sigmoid function scales each value into the range [0, 1], which is treated as an independent weight score for each channel; and the previous convolutional layer’s weights multiply the weight score as the re-scored weights.

The four Conv blocks increase the input channel number from 1 to 256. A global max-pooling layer is applied to the last Conv block to extract each channel’s max value. A flatten layer combined these features and reshaped the size to *b* × 256. The following four dense blocks reduce the feature size from *b* × 256 to *b* × 1 to get the predicted GDT-TS score.

Except the sigmoid activation function used in the two SE blocks and the ReLu activation used in the last output layer, all the other layers use the Leakey-ReLu activation function if applicable. The deep learning network above was trained with the Smooth L1 loss function [22]. Unlike the mean squared error (MSE) loss and standard L1-loss function, the Smooth L1 loss is less sensitive to the outliers and derivable at 0 point. The **Equation 1** is the formula of the smooth L1 loss, where *x* denotes the difference between the predicted and true GDT-TS scores. It is a combination of MSE loss and L1 loss. The derivative of the smooth L1 loss is represented by **Equation 2**. The derivative is x when x is in the range [1, 1], which is linear. Otherwise, it is a constant (1 or −1). This property ensures the deep network is stable and converges fast.

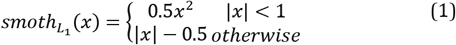

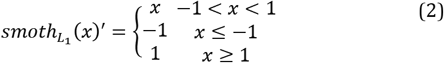

For the convolutional and linear layers, we utilized the kaiming initialization [23] to initialize the start values. We implemented the deep network with PyTorch [24]. It was trained by Adam optimization method [25] with *β*_1_ = 0.9 and *β*_2_ = 0.999. The learning rate was set as a constant value of 0.00005 and the batch size as 16.

### Datasets and evaluation metrics

We generated the difference map for each structural model predicted for CASP8-13 targets by CASP8-13 structure prediction servers. Each CASP protein target may have up to a few hundred structural models (decoys). The true GDT-TS scores of these models were calculated as labels to train the deep learning method. 120,064 structural models were used for training and validation, and 14,580 structural models of CASP13 were used as the test dataset. CASP8-12 targets used for training have different sequence lengths (see **Figure 3** for the length distribution). To improve the effectiveness of training on the models of different lengths, the CASP8-12 models were divided into many batches, each of which consisted of 16 structural models of the same length. 80% of randomly selected models in each batch were pooled together to form the training dataset and the remaining models were used as the validation dataset.

**Figure 3.**
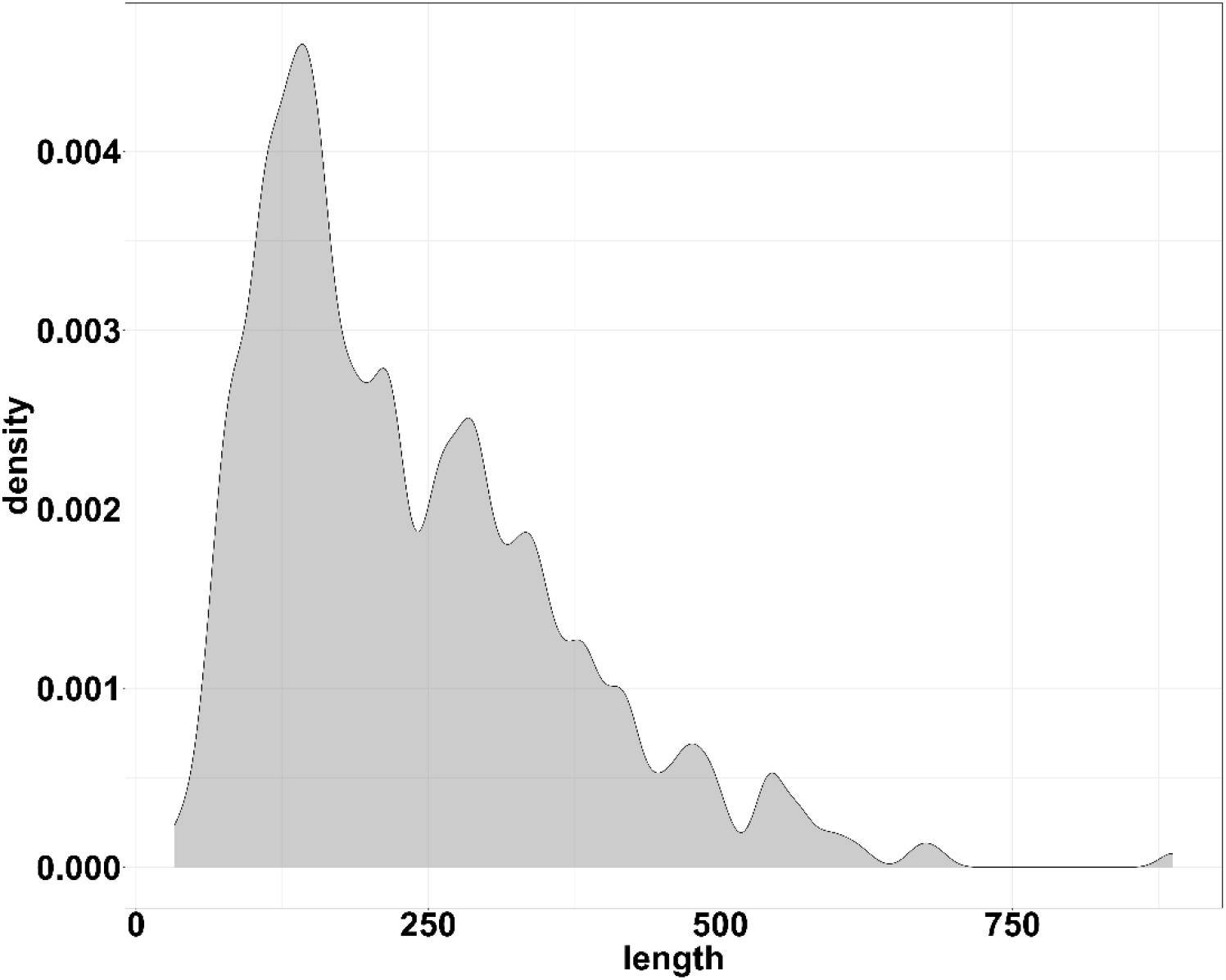
The distribution of the sequence length of CASP8-12 targets.

The predicted performance of DISTEMA and other methods for a target was evaluated by the GDT-TS score loss of ranking the models of the target, which is defined as the absolute difference between the true GDT-TS score of the best model of a target and that of the top model selected by the predicted GDT-TS scores of the models of the target. A ranking loss of 0 means that the best model for a target has been selected by the predicted GDT-TS scores. The average GDT-TS loss of ranking models of all the targets in the test dataset was used to evaluate the performance of the EMA methods. Moreover, the Pearson’s correlation coefficient between the predicted GDT-TS scores of the models of a target and their GDT-TS scores was calculated. The average Pearson’s correlation coefficient over all the targets in the test dataset was also employed to estimate the performance of the EMA methods.

## Results and Discussion

Results on the CASP13 dataset and comparison with single-model QA methods without using inter-residue distances.

We evaluated DISTEMA with several single-model EMA methods on CASP13 dataset. The results of ProQ2 [26], ProQ3 [9], ProQ3D [10], ProQ4 [11], and two Oromia methods [27] on the CASP13 dataset reported in [8] were compared with that of DISTEMA. The average ranking loss and Pearson’s Correlation Coefficient (PCC) of these methods are reported in **Table 1**. DISTEMA and ProQ4 have the lowest ranking loss of 0.079, which is a 9% improvement over the second-lowest loss of 0.086. The PCC of DISTEMA is 0.831, 19% higher than the second highest PCC achieved by ProQ4.

**Table 1.** Seven models per-target comparison on CASP13. DISTEMA gets the TOP first-ranked loss and PCC performance.

| Model | first-ranked loss | PCC |
| --- | --- | --- |
| DISTEMA | **0.079** | **0.929** |
| ProQ4 | **0.079** | 0.711 |
| ProQ3D | 0.086 | 0.652 |
| ProQ3 | 0.089 | 0.594 |
| Koroma-B | 0.095 | 0.095 |
| VoroMQA-A | 0.109 | 0.642 |
| ProQ2 | 0.114 | 0.604 |

**Figure 4** is the scatter plot of the true GDT-TS score of the best model of each target against the true GDT-TS score of the top model selected by DISTEMA for the target. The solid red line denotes the regression line between predicted GDT TS and true GDT TS and the yellow line is the 45-degree line on which points have 0 loss. Larger the distance between a data point and the yellow line, bigger the loss is. For four targets (i.e., T0949, T0987, T0980s2, T1019s2), their best models were successfully selected as top models by DISTEMA, yielding a loss of 0. Two outliers - T1008, T1022s2 – have the largest loss for DISTEMA. **Figure 5** illustrates the distribution of the ranking losses of DISTEMA on the CASP13 dataset. The vertical dashed black line is the mark for the mean loss. More data points are located on the left side of the black line. The skewness of the distribution 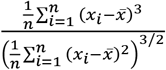 is 2.377.

**Figure 4.**
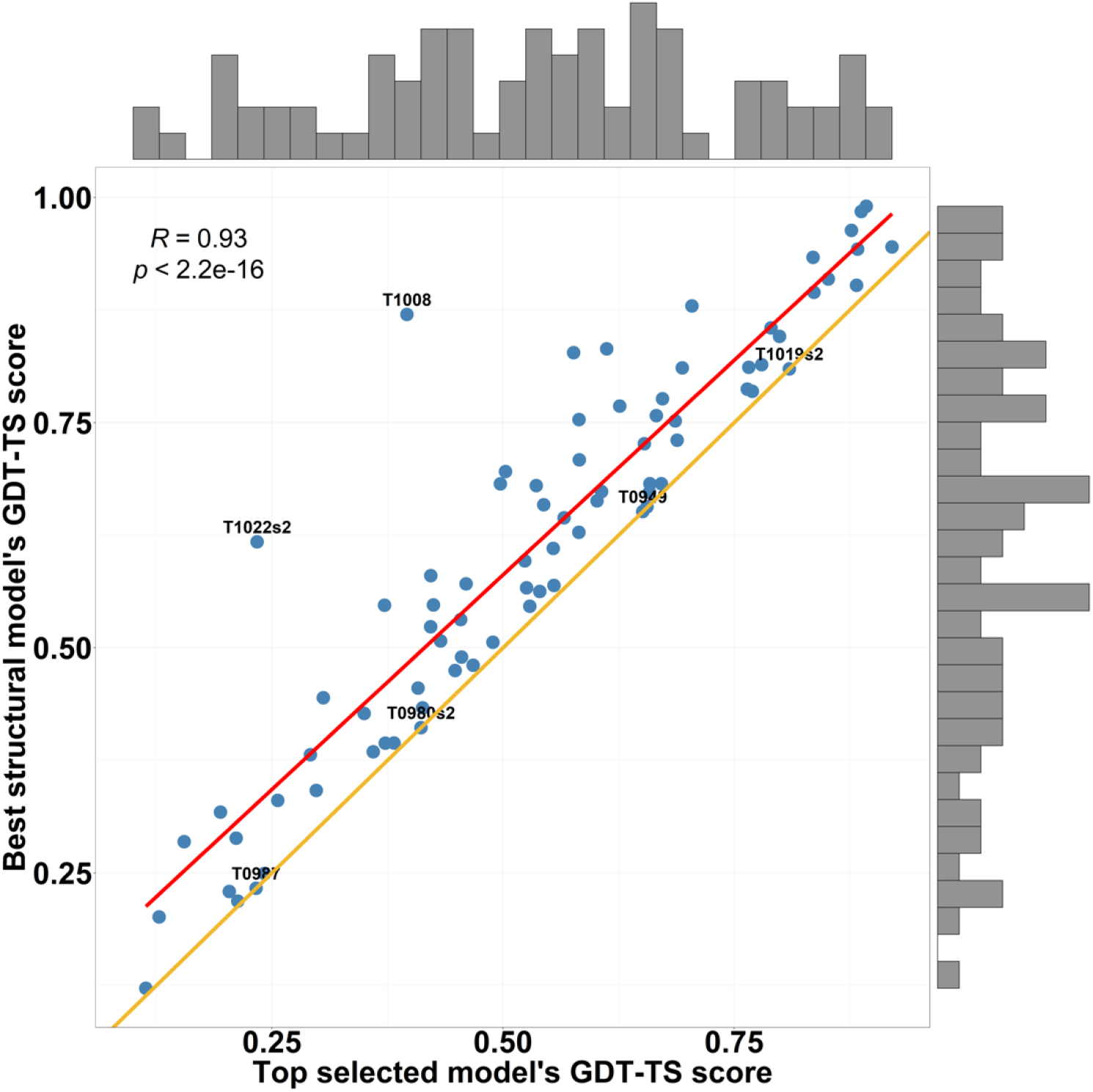
The plot of the true GDT-TS scores of the best models against the true GDT-TS scores of the top models selected by DISTEMA for 80 CASP13 targets. The histogram at the top is the distribution of the GDT-TS scores of the top selected models. The histogram on the right shows the distribution of the GDT-TS scores of the best models. The yellow is a 45-degree line with the slop of 1. The points on the yellow line represent the targets whose best models and top selected models have the same GDT-TS score (i.e., 0 loss). Four targets (T0949, T0987, T0980s2, T1019s2) have 0 loss. Closer a point to the yellow line, lower loss of GDT-TS score for the target. Two targets (T1008 and T2022s2) are the two outliers with very high loss. The red line is the linear regression line between the two groups of GDT-TS scores. The correlation between the two groups of the scores is 0.93.

**Figure 5.**
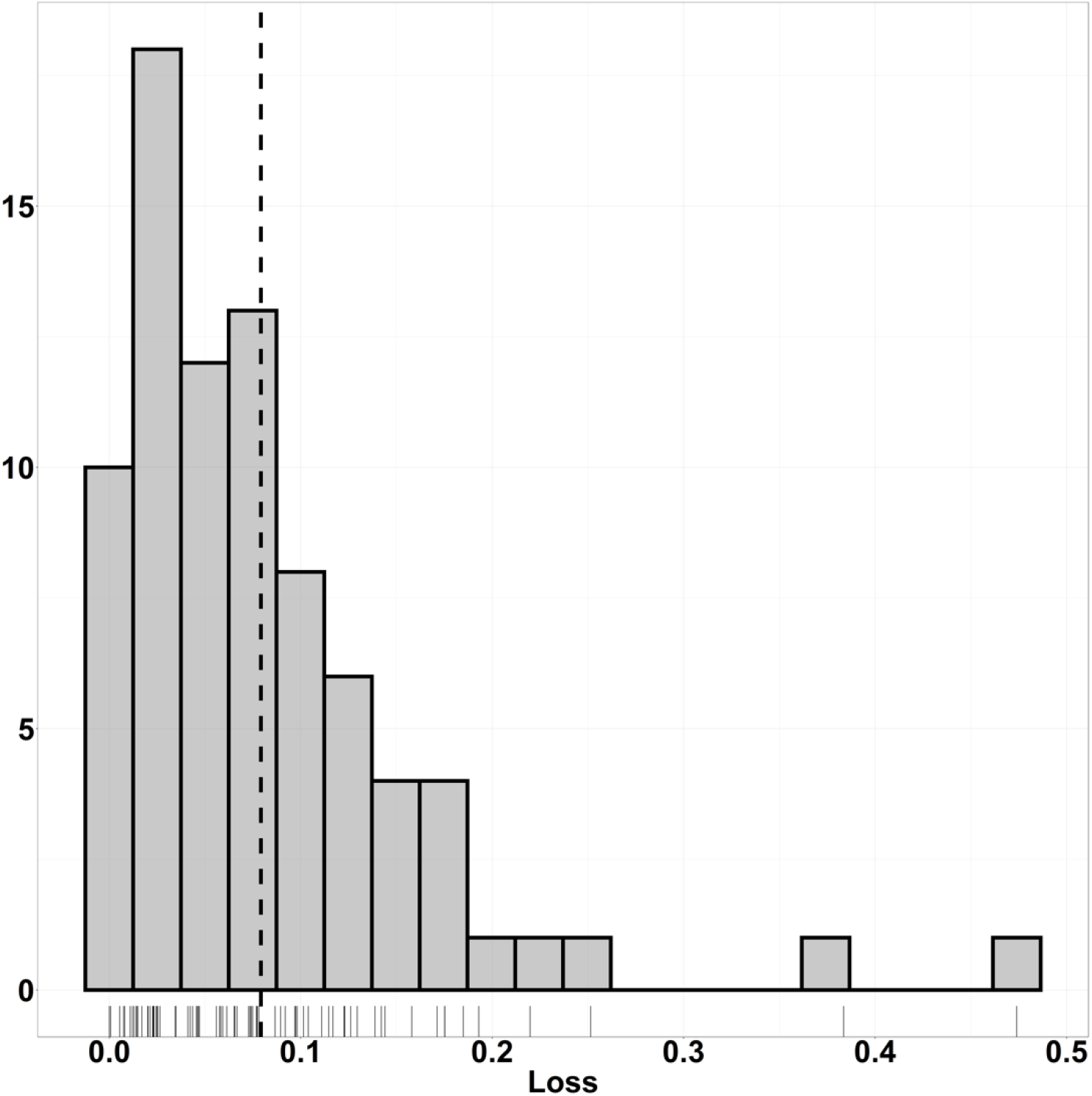
Histogram of the distribution of the ranking loss of DISTEMA over the 80 CASP13 targets. The vertical black dashed line represents the position of the mean value.

In addition to the ranking loss, we applied a non-parametric method Kolmogorov Smirnov test (KS test) to measure the distance between the distribution of true GDT-TS scores and that of predicted GDT-TS scores of the CASP13 models. We conducted the KS test on the two datasets. The first dataset contains the true GDT-TS scores of all the CASP13 models, and their GDT-TS scores predicted by DISTEMA. The distributions of the two kinds of scores were compared. The second dataset include the true GDT-TS scores of the best modes for the CASP13 targets and the true GDT-TS scores of the top models selected for them. For both tests, we used the same null hypothesis *H0*: *no difference between the two distributions*. We calculated Kolmogorov–Smirnov statistics *D*_(*n,m*)_ (i.e., 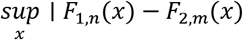) the measurement of the difference between the two distributions. Here, *sup_x_* is the supremum function, which in this case is considered as the *max* function. *F*_1,*n*_ and *F*_2,*m*_ are the empirical distribution functions for first and second sample respectively, where *n* and *m* are sample size.

On the dataset 1, D-statistics of the KS test is 0.11266, and the p-value (2.2e – 16) is smaller than a significance threshold (i.e., 0.05), which means these two samples come from different distributions. **Figure 6** shows the two samples’ cumulative distribution function (CDF) curves. The red vertical dashed line is the D-statistics, representing the maximum absolute difference between these two CDF. The shapes of the two curves are different.

**Figure 6.**
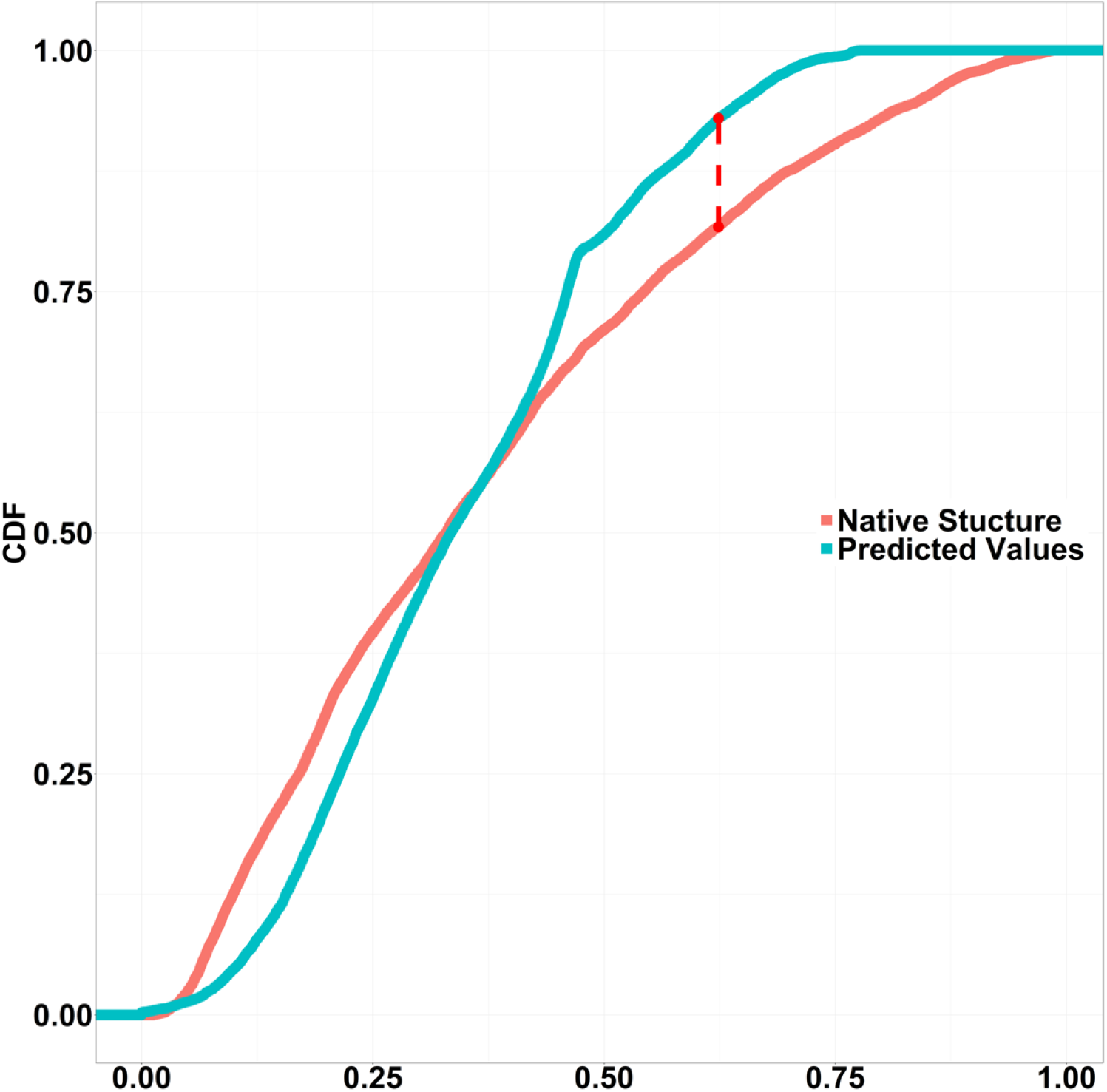
K-S plot for global GDT TS V.S. predicted GDT TS. D statistics: 0.11266, p-value:2.2e-16.

The analysis shows that the distribution of the true GDT-TS scores of all the models and the distribution of their predicted GDT-TS scores have somewhat different distributions, indicating that the quality scores of some models (e.g., some models of very low quality) are hard to predict. Enlarging the training dataset may alleviate the problem.

In contrast, on the dataset 2, the D-statistics is 0.2 and the p-value of KS-test is 0.08152, higher than the threshold, suggesting the null hypothesis be accepted. That is, the distribution of the true GDT-TS score of the best model for each target has no difference than the distribution of the true GDT-TS score of the top model selected for each target. **Figure 7** illustrates the two distributions’ CDF curves, where the blue line and red line are generally in the same shape. The similar distribution of the GDT-TS scores of best models and top selected models further confirm that the ranking capability of DISTEMA is sound.

**Figure 7.**
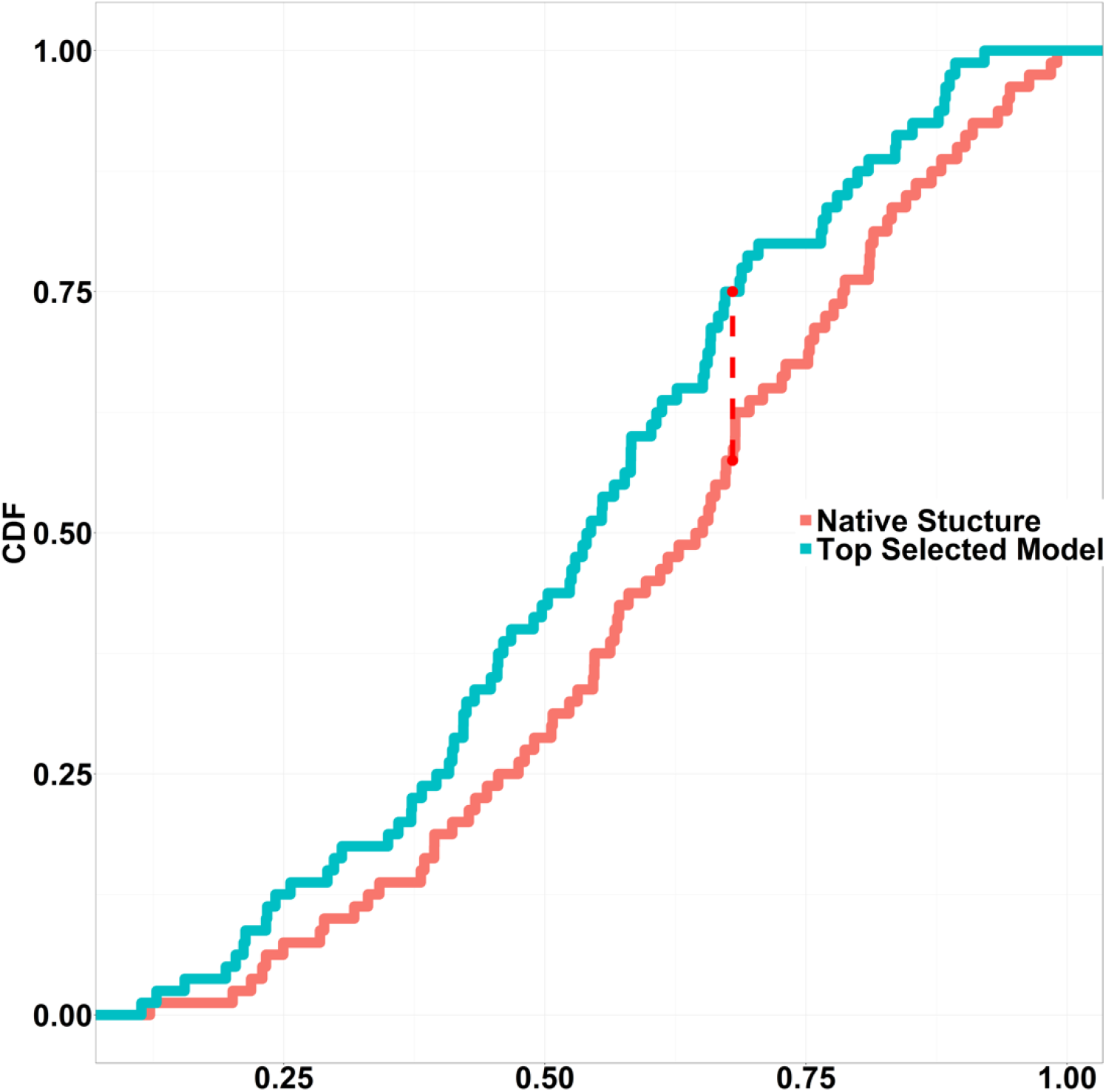
K-S plot for per target GDT TS V.S. predicted GDT TS. D statistics:0.2, p-value:0.08152.

### Comparison with a distanced-based EMA method

We compared DISTEMA with a recent distance-based single-model method QDeep [28]. QDeep uses one dimensional CNN with inter-residue distance features derived from distance predictions, sequence information, and energy scores, while DISTEMA only uses the raw distance maps as input. DISTEMA was evaluated on the same 3000 models of 20 CASP13 targets on which QDeep was evaluated. **Table 2** reports the results of the two methods. DISTEMA performed better than QDeep according to the ranking loss even though it only used one kind of input information, but worse than QDeep according to Pearson’s correlation. The results show that using only raw distance maps with deep learning can predict the quality of a single protein model reasonably well and integrating other features with the distance information may further improve the prediction performance.

**Table 2.** The ranking loss and Pearson’s correlation coefficient of DISTEMA and QDeep.

| Methods | Ranking Loss | Pearson's Correlation |
| --- | --- | --- |
| QDeep | 0.088 | **0.866** |
| DISTEMA | **0.083** | 0.826 |

### Contribution of Squeeze-and-Excitation (SE) blocks with attention

We trained two deep learning networks to investigate the impact of SE blocks with attention. The two networks have the same architecture except one network has SE blocks, but another does not. The two networks were trained with the same experimental setting and were evaluated on the CASP13 dataset. The network with SE blocks has the ranking loss of 0.079, 7.5% lower than 0.085 of the network without SE blocks, indicating that attentive SE blocks can improve the performance of model quality prediction. The attention mechanism in SE blocks can more effectively pick up the relevant features anywhere in the input and assign them higher weights to improve the prediction performance.

## Conclusion and Future work

We designed and developed an attentive 2D CNN with the channel-wise attention to directly leverage a raw inter-residue distance map to predict the global quality of a single protein model. Using only the protein distance information, the deep learning method with the attention mechanism is able to automatically extract features relevant to model quality from the raw input and achieves the lower model ranking loss than other state-of-the-art single-model EMA methods that use various expert-curated protein structural features. The results demonstrate that raw protein distance maps contain substantial information that can be captured by advanced deep learning methods to estimate the accuracy of a single protein model. In the future, larger training datasets, additional input features, and more advanced deep learning architectures [29] can be used to further improve the distance map-based methods for improving the prediction of protein model quality.

## Acknowledgements

We thank CASP for providing the data for public use.

## Availability of data and materials

Availability: https://github.com/jianlin-cheng/DISTEMA

## Authors’ contributions

Jianlin Cheng and Xiao Chen designed this project. XC implemented it and collected the results. XC and JC wrote the manuscript.

